# *pumilio* regulates sleep homeostasis in response to chronic sleep deprivation in *Drosophila melanogaster*

**DOI:** 10.1101/833822

**Authors:** Luis A. De Jesús-Olmo, Norma Rodríguez, Marcelo Francia, Jonathan Alemán-Rios, Carlos J. Pacheco-Agosto, Joselyn Ortega-Torres, Richard Nieves, Nicolás Fuenzalida-Uribe, Alfredo Ghezzi, José L. Agosto

**Affiliations:** Department of Biology, University of Puerto Rico, Rio Piedras, P.R., USA

**Keywords:** sleep homeostasis, neuronal homeostasis, Pumilio, pum, Drosophila, chronic sleep deprivation, synaptic proteins, neuronal excitability

## Abstract

Recent studies have identified the *Drosophila* brain circuits involved in the sleep/wake switch and have pointed to the modulation of neuronal excitability as one of the underlying mechanisms triggering sleep need. In this study we aimed to explore the link between the homeostatic regulation of neuronal excitability and sleep behavior in the circadian circuit. For this purpose, we selected the neuronal homeostasis protein Pumilio (Pum), whose main function is to repress protein translation and has been linked to modulation of neuronal excitability during chronic patterns of altered neuronal activity. Here we explore the effects of Pum on sleep homeostasis in *Drosophila melanogaster*, which shares most of the major features of mammalian sleep homeostasis. Our evidence indicates that Pum is necessary for sleep rebound and that its effect is more pronounced during chronic sleep deprivation (84 hours) than acute deprivation (12 hours). Knockdown of *pum*, results in a reduction of sleep rebound during acute sleep deprivation and the complete abolishment of sleep rebound during chronic sleep deprivation. These behavioral changes were associated with accompanying changes in the expression of genes involved in the regulation of neuronal excitability. Interestingly, *pum* knockdown also increased baseline daytime sleep, suggesting that Pum differentially regulates rebound and normal sleep. Based on these findings, we propose that Pum is a critical regulator of sleep homeostasis through neural adaptations triggered during sleep deprivation and induces rebound sleep by altering neuronal excitability.

## 1 Introduction

It is well established, even by our own experience, that the urge to sleep increases as a function of time awake. This urge, or sleep drive, triggers a prolonged compensatory sleep after the organism is sleep deprived (Daan et al., 1984; Allada, et al., 2017). This compensatory sleep, which is also called sleep rebound, is a key indicator of the homeostatic regulation of sleep (Vyazovskiy, et al., 2009). In this process, deviations from a reference level of sleep are compensated, i.e. lack of sleep fosters compensatory increase in the intensity and duration of sleep, whereas excessive sleep counteracts the sleep need (Tobler and Ackermann, 2007). More than a century of sleep research has made important progress in understanding the function of sleep and its regulatory circuitry, but the molecular basis of sleep homeostasis remains elusive (Cirelli & Tononi, 2008; Siegel 2008; Sehgal et al., 2007; Donlea 2017). Understanding the molecular mechanisms involved in the regulation of sleep homeostasis is key for the overall understanding the regulation of both the sleep circuit and the sleep function. To achieve that level of understanding, we need to study the link between molecular markers, sleep brain circuits and homeostatic sleep behavior.

The fruit fly *Drosophila melanogaster* is an ideal model to study the molecular markers impacting sleep behavior. Sleep rebound is a stable phenotype in flies which shares most major features of mammalian sleep homeostasis (Huber, et al., 2004). *Drosophila* shows easily measurable and recognizable sleep patterns linked to reduced brain activity (Nitz et al., 2002; Van Swinderen et al., 2004), limited sensory responsiveness during sleep and display a robust homeostatic sleep rebound (Hendricks et al., 2000; Shaw, et al., 2000) as occurs in mammals. Moreover, it has been demonstrated that humans and fruit flies have a common sleep control mechanism involving GABA receptors in brain neurons linked to the circadian clock (Parisky, et al., 2009; Chung, et al., 2009). In addition, fly genetics has been used as a tool to validate human sleep biomarkers affected by sleep deprivation (Thimgan et al., 2013). Hence, we circumscribed our study of the molecular relationship between homeostatic markers and sleep behavior to the fly model.

Recent studies have shown that two structures of *Drosophila’*s brain central complex, the Ellipsoid Body (EB) and the fan body (FB), induce sleep when artificially activated, and produce insomnia, when inhibited (Liu, et al., 2016; Donlea, et al., 2011). Other studies have shown that neuronal microcircuits in the mushroom body (MB) drives rebound recovery after sleep deprivation (Sitaraman, et al., 2015). Follow up studies have produced important progress by identifying dopamine as the neuromodulator responsible for the homeostatic switch operation between sleep/wake, which is mediated by potassium currents (Pimentel, et al., 2016). Homeostatic sleep seems to be controlled by the dorsal FB neurons, which are electrically active during wake and electrically silent during rest (Pimentel, et al., 2016). These studies point to the regulation of neuronal excitability as an important effector of the sleep regulation. Nevertheless, the underlying molecular framework that connects neuronal excitability with sleep behavior is a relatively unexplored area of research.

Several genes have been identified to regulate normal sleep, but only a few genes have been linked to the molecular regulation of homeostatic sleep compensation after sleep deprivation. A mutation in the *Shaker* (*Sh*) gene, which encodes a voltage dependent potassium channel involved in membrane repolarization, increases neuronal excitability and reduces normal sleep (Cirelli et al., 2005), but fails to alter sleep rebound. Interestingly, the *Shaker* activator *sleepless* (*sss*), which encodes for a brain-enriched glycosyl-phosphatidylinositol-anchored protein, impairs sleep rebound (Koh, et al., 2008), perhaps by a mechanism independent of *Shaker*. The gene *crossveinless* (*cv-c*), which codes for a Rho-GTPase-activating protein, is necessary for dorsal FB neurons to transduce the excitability produced by sleep pressure into homeostatic sleep (Donlea, et al., 2014). Knocking down the *Cullin 3* (*Cul3*) ubiquitin ligase gene and its putative adaptor *insomniac (inc)*, reduces sleep rebound after sleep deprivation (Pfeiffenberger & Allada, 2012). Mutants of fragile X mental retardation gene (*Fmr1*), a translational inhibitor that causes the most common form of inherited mental retardation in humans, have also been reported to reduce sleep rebound (Bushey, et al., 2009). In addition, it was reported that interfering with the expression of the genes *sandman* (*sand*) and *Sh* in the dorsal FB neurons, increased or decreased sleep respectively as part of the sleep/wake switch (Pimentel, et al., 2016). The regulatory picture presented by these genes and the other neuromodulators and proteins known to affect homeostatic sleep compensation seems far from complete, although together, they also point to neuronal excitability as a key component of sleep homeostatic regulation.

Unregulated neuronal excitability may lead to a potentially disruptive positive feedback. To cope with this, neurons have evolved compensatory mechanisms to reduce excitability. The mechanisms by which neurons stabilize firing activity have been collectively termed “homeostatic plasticity” (Marder & Prinz, 2003; Turrigiano & Nelson, 2004; Turrigiano 2008; 2012; Davis 2006; Pozo & Goda, 2010). Therefore, it is plausible that wake promoting neurons, after prolonged times of wakefulness, would utilize one of the homeostatic plasticity mechanisms to regulate neuronal excitability. In this study, we begin to explore the relationship between neuronal homeostasis mechanisms and sleep regulation by testing the role of the neuronal homeostasis gene *pumilio* (*pum*) on the regulation of compensatory sleep.

The protein encoded by *pum* is characterized by a highly conserved RNA-binding domain, which acts as a post-transcriptional repressor of mRNA targets. Binding occurs through an RNA consensus sequence in the 3’-UTR of target transcripts—the Pumilio Response Element (PRE), 5’-UGUANAUA-3’, that is related to the Nanos Response Element (NRE) (Wang et al., 2018). While it was originally described in *Drosophila* for its critical role in embryonic development, Pum has an important role in the development of the nervous system. Pum is known for controlling the elaboration of dendritic branches (Ye, et al., 2014), and is also required for proper adaptive responses and memory storage (Dubnau, et al., 2003). Evidence of its regulatory role if neuronal homeostatic processes include Pum’s repression of translation of the *Drosophila* voltage-gated sodium channel (*paralytic*) in an activity dependent manner (Mee, et al., 2004; Murano, et al., 2008). Pum-mediated repression of the voltage gated sodium channel plays a pivotal role in the regulation of neuronal homeostasis, given the central role of the sodium channel in the regulation of membrane excitability (Weston & Baines, 2007). Furthermore, *pum* was found to be necessary for the homeostatic compensation of increased neuronal activity, or what is known as homeostatic synaptic depression (Fiore, et al., 2014). In addition, Pum has been found to influence synaptic bouton size/number, synaptic growth and function by regulating expression of eukaryotic initiation factor 4E (eIF4E), which is the limiting factor for the initiation of the CAP dependent translation in Eukaryotes (Menon, et al., 2004; Vessey, et al., 2006; Cao, et al., 2009). Pum was our first choice to study neuronal homeostasis effects on compensatory sleep because microarray experiments show that *pum* is expressed in PDF-expressing cells, which are key circadian cells known to promote wakefulness in *Drosophila* (Kula-Eversole, et al., 2010; Parisky, et al. 2008). With over 1000 potential targets and many others indirect targets through its eIF4E regulatory role, based on the cumulative evidence, Pum could be considered a master regulator of neuronal homeostatic processes (Gerber, et al. 2006; Chen, et al. 2008; Menon, et al. 2004).

Our data shows that sleep rebound is reduced by knocking down *pum* in the circadian circuit. This effect is more pronounced after chronic sleep deprivation in comparison with acute sleep deprivation. Our behavioral and molecular data correlates with *pum’s* differential involvement in regulating compensatory sleep as a function of sleep need. This, in turn, suggests a mechanistic framework for linking sleep function and regulation through neuronal homeostasis mechanisms.

## Results

### *Pumilio* regulates sleep rebound differentially between acute and chronic mechanical sleep deprivation

Studies exploring the mechanisms of neuronal homeostasis often involve long-term manipulations of neural activity, spanning from 48 hours to the entire life span (Davis, 2013; Turrigiano et al., 1998; Turrigiano, 2012). Moreover, studies linking *pum* with neuronal homeostasis primarily use genetic manipulations that alter neuronal activity throughout the lifetime of the organisms (Weston and Baines, 2007; Mee et al., 2004; Muraro et al., 2008). Thus, in this study we decided to explore the role of *pum* in the regulation of sleep homeostasis induced by chronic (long-term) sleep deprivation as well as acute sleep deprivation (SD).

We knocked down the expression of *pum* using a transgenic fly containing a *pum* RNA interference construct (*pum*^RNAi^) under control of the upstream activating sequence (UAS) of the yeast transcription factor Gal4. To activate the UAS-*pum*^RNAi^ we used a second transgenic construct that expressed Gal4 under control of the *timeless* (*tim*) gene promoter (*tim-*Gal4). When both transgenes are present in the same fly (*tim-*Gal4/UAS-*pum*^RNAi^), the *pum*^RNAi^ construct is expressed constitutively in *tim* expressing neurons. We selected the *tim*-Gal4 driver because it is a strong and broadly expressed promoter targeting circadian cells found in several brain structures including the wake promoting, PDF-expressing ventral lateral neurons and both the EB and FB neurons (Kaneko & Hall 2000).

In our first set of experiments, we subjected the pum^*RNAi*^ (UAS-*pum*^RNAi^/*tim-*Gal4) and their “sibling” control flies (UAS-*pum*^RNAi^/+), which carry the *pum*^RNAi^ construct by itself, to either chronic or acute mechanical SD protocol. In both protocols, flies were placed in the Drosophila Activity Monitors to be monitored for 6 days for baseline sleep. After the 6th day, flies were subjected to mechanical SD using an apparent random shaking program (see methods). Both chronic and acute deprivation protocols were identical in terms of stimulus intensity and pattern; the only difference was the duration of the deprivation period. For chronic sleep deprivation, the SD protocol was active for the first 84 hours starting at the beginning of the first dark period (Fig.1), while for acute sleep deprivation, the SD protocol lasted only 12 hours, which encompassed the entirety of the dark period preceding the sleep recovery period.

The results from the chronic SD showed a strong effectiveness of the sleep deprivation method during the first 12 hours (Fig. 1A). However, as time progressed, we noticed a gradual increase in the amount of sleep in all the sleep deprived genotypes during sustained mechanical deprivation. However, this increase in sleep through time did not seem to affect the sleep rebound, as control flies were able to produce a normal sleep rebound pattern that initiated at the 84^th^ hour—immediately after the SD protocol was terminated (Fig. 1 A-B). Surprisingly, we noticed that *pum*^RNAi^ flies did not show any rebound (Fig. 1C). To determine if this lack of sleep rebound was related to an insufficient sleep deprivation, we quantified the sleep lost and used this value to normalize the sleep recovery after deprivation. The quantification of cumulative sleep loss during the 84-hour deprivation period showed a significant difference between the *pum*^RNAi^/*tim-*Gal4 flies and the *tim-*Gal4/+ control flies, but no difference between the *pum*^RNAi^/*tim-*Gal4 flies and the UAS-*pum*^RNAi^/*+* controls (Fig 1D). The fact that this difference was not significant between both controls and pum^*RNAi*^ flies, suggests the difference in effectiveness could be due to the genetic background rather than the knockdown of *pum*. The results for sleep recovery show a normal recovery pattern in both controls after sleep deprivation as indicated by the increase in cumulative sleep recovered during the first hours after SD, when compared to non-sleep deprived flies during the same time period (Fig. 1E). After normalizing by the sleep lost, *pum*^RNAi^ flies showed a negative sleep recovery, which indicates *pum*^RNAi^ flies were more active than the non-deprived controls after 84 hrs of continuous deprivation (Fig. 1E). This loss of homeostatic regulation in the recovery of *pum*^RNAi^ flies was maintained up to 96 hours post-deprivation (see supplementary figure S2). In our experiments, the UAS-*pum*^RNAi^/*+* control lines are siblings of the UAS-*pum*^RNAi^/*tim-*Gal4 flies. Meanwhile the *tim-*Gal4/+ controls were generated directly by crossing the parental *tim*-Gal4 line with a non-transgenic wild-type (CS), which can introduce differences in genetic background. Thus, our conclusions are based mostly on the results from “sibling controls” because they have a greater genetic similarity, which results in a more similar baseline sleep pattern than parental controls (Figs. 1 A-C). Hence, for the acute SD experiments, parental controls were not used.

**Figure 1:**
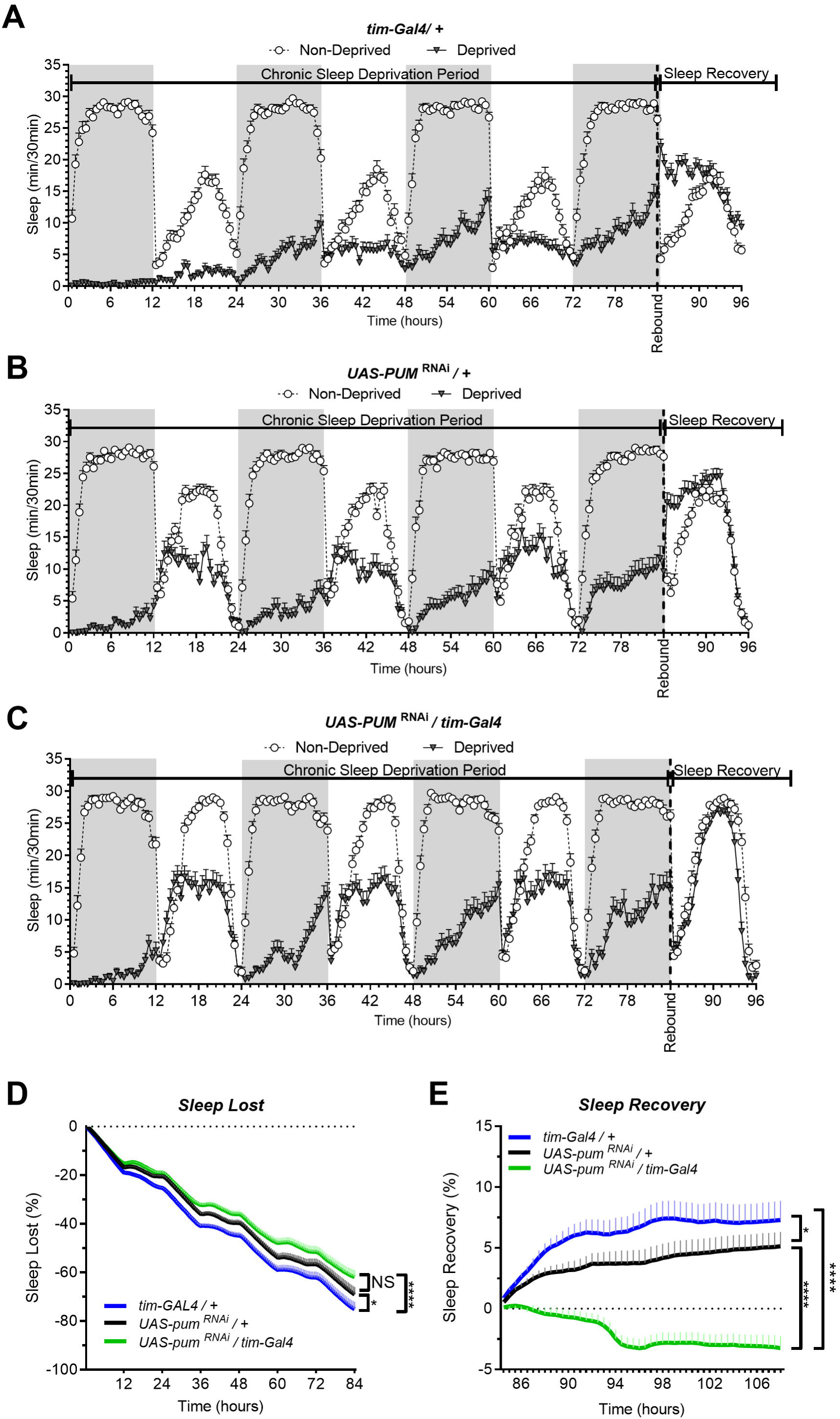
*Pum* knockdown eliminates sleep recovery after chronic mechanical sleep deprivation. Sleep comparison of UAS*-pum*^RNAi^*/tim-*Gal4 (experimental) vs *tim-*Gal4*/+* (parental) flies and UAS-*pum*^RNAi^*/+* (“sibling” controls) during chronic SD. The X axis indicates time after start of sleep deprivation. **(A-C)** Depiction of sleep activity during the sleep deprivation and sleep rebound period for all genotypes. **(D)** Cumulative sleep lost during deprivation expressed as a percentage of total sleep in non-deprived flies of the same genotype. Two-way ANOVA using “genotype” as a factor and “time” as a repeated measure showed significant differences in genotypes (F (2, 132) = 11.24 P<0.0001), time (F (167, 22044) = 1033 P<0.0001) and interaction (F(334, 22044) = 3.066, P<0.0001). Post-hoc analysis using Dunnett’s multiple comparisons test showed significant differences between *UAS-pum*^RNAi^*/tim-*Gal4 *vs tim-*Gal4*/+* flies (P<0.0001). **(E)** Percent sleep recovery after SD. Two-Way ANOVA with repeat measures indicated significant differences in genotypes (F (2, 132) = 18.58 P<0.0001) and interaction (F (94, 6204) = 13.73 P<0.0001). Post-hoc analysis using Sidak’s multiple comparisons test comparing both control genotypes against experimental flies, revealed significant differences (P<0.0001) between UAS*-pum*^RNAi^*/tim-*Gal4 *vs tim-*Gal4*/+* flies and UAS-*pum*^RNAi^*/+* throughout the recovery period. The data shown represents two experiments with the following sample sizes (N): *tim-*Gal4*/+* Non-Deprived (N=56) and Deprived (N=53); UAS-*pum*^RNAi^*/+* Non-Deprived (N=60) and Deprived (N=35); UAS-*pum*^RNAi^*/+* Non-Deprived (N=63) and Deprived (N=39). Because the calculations of sleep lost and sleep recovery involve both the Non-Deprived and Deprived groups (see methods), the N for panels A and B is equal to the N of the Deprived group. SD. Data points and error bars represent means ± SEM. Stars indicate significance level (* denotes p<0.05; ** p< 0.01; *** p< 0.001; **** p< 0.0001).

The results from the 12 hours acute SD showed sleep lost effectivity close to 100% for both *pum*^RNAi^ and “sibling” controls (Fig. 2A-B). During the deprivation period (0 to 12 hours), the cumulative sleep loss in deprived flies did not show a significant difference between the two genotypes (Fig. 2E) Once again, controls showed an effective sleep rebound (Fig. 2A), while *pum*^RNAi^ flies showed a reduction in sleep rebound (Fig. 2B). However, this time the rebound was not completely abolished as we observed during chronic SD (Fig. 2B vs 2D). We included the chronic deprivation rebound period as a point of comparison between acute vs chronic (Figs. 2C-D). The results from the acute SD sleep recovery resembled the results from chronic SD with a normal rebound in “sibling” controls and reduced sleep recovery in *pum*^RNAi^ flies. Nevertheless, the sleep recovery of *pum*^RNAi^ flies was not negative as we observed during chronic SD (Fig. 2F). When acute vs chronic SD results are compared (Fig 2G), we see significant differences, not only between the genotypes, but also within *pum*^RNAi^ flies exposed to acute vs chronic SD, while the rebound difference of the “sibling” control between acute vs chronic SD remains constant. These results suggest that *pum* differentially regulates acute vs chronic SD. This interpretation is in fact reinforced by our molecular experiments contrasting gene expression changes between acute and chronic SD as reported below and in the supplementary material (supplementary Fig S3).

**Figure 2:**
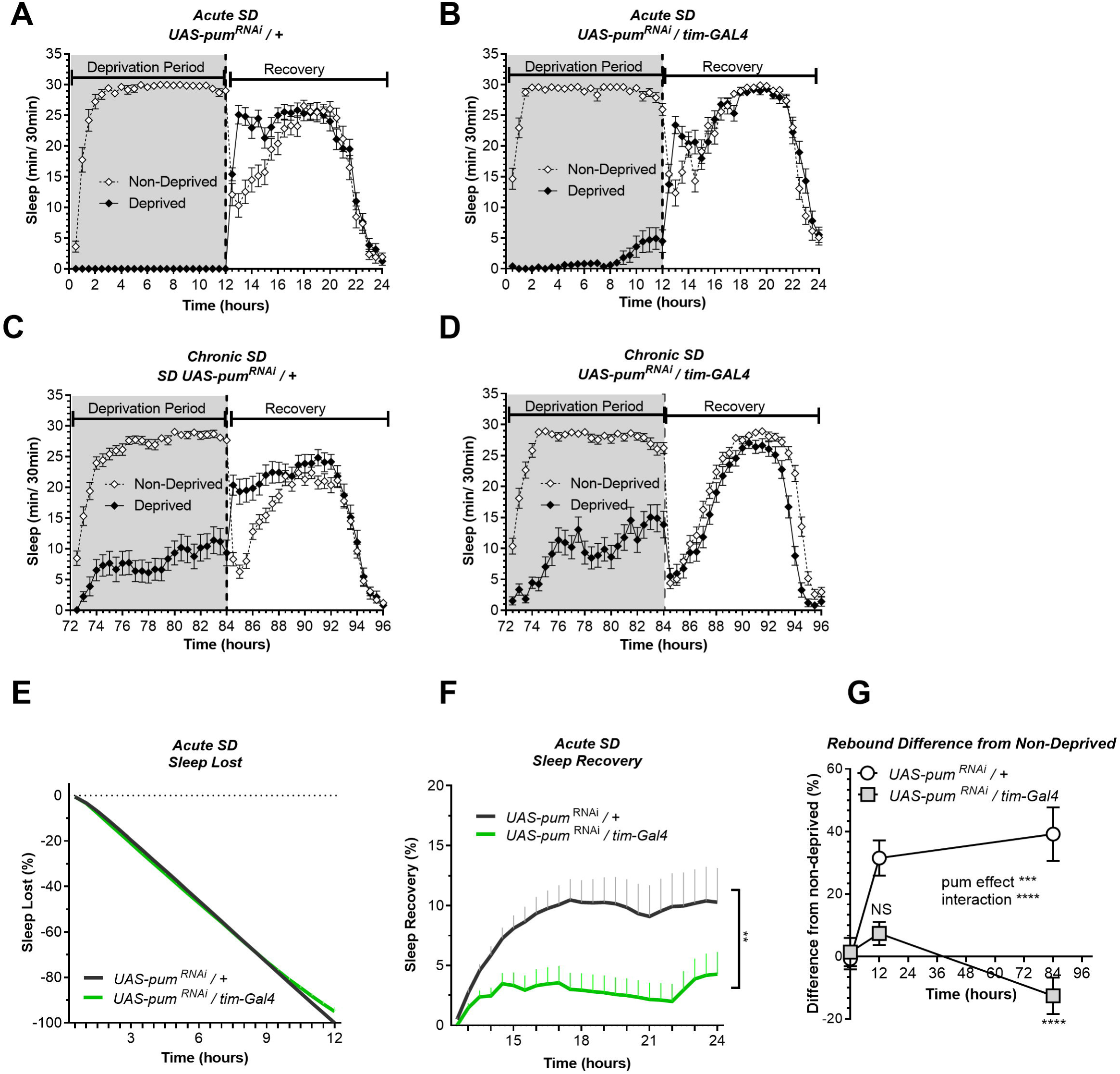
*Pum* knockdown differentially reduces sleep recovery in acute vs chronic SD. Sleep comparison of UAS*-pum*^RNAi^*/tim-*Gal4 (experimental) vs UAS-*pum*^RNAi^*/+* (“sibling” controls) during acute (12 hours) mechanical SD. The X axis indicates time after sleep deprivation. **(A-B)** Depiction of sleep activity during the sleep deprivation and sleep rebound period for both genotypes during acute SD. **(C-D)** Depiction of the sleep activity during sleep deprivation and sleep rebound period for both genotypes during hours 72 to 96 of chronic mechanical SD included for ease of comparison. The y-axis shows the number of minutes that flies slept in intervals of 30 min. **(E)** Cumulative sleep lost during deprivation expressed as a percentage of total sleep in non-deprived flies of the same genotype. Two-way ANOVA, using “genotype” as a factor and “time” as a repeated measure, did not showed significant differences between the genotypes (P=0.8664). **(F)** Percent sleep recovery after SD. Two-Way ANOVA with repeat measures indicated significant difference in genotypes (F (1, 58) = 7.114, P<0.0099) and interaction (F (23, 1334) = 3.054, P<0.0001). **(G)** Percent difference in rebound between deprived and non-deprived flies after acute and chronic sleep deprivation protocols of UAS*-pum*^RNAi^*/+* and UAS*-pum*^RNAi^*/tim-*Gal4 flies. Two-way ANOVA with repeated measures showed a significant difference in genotype (F (1, 91) = 13.72, P=0.0004) and time vs genotype interaction (F (2, 106) = 13.97, P<0.0001). Post-hoc analysis using Tukey’s multiple comparisons test revealed significant differences between UAS*-pum*^RNAi^*/+* and UAS*-pum*^RNAi^*/tim-*Gal4 at 84 hours of deprivation (P<0.0001) no difference was observed at 12 hours (acute SD) (P=0.0735). The data shown represents one experiment with the following sample sizes (N): UAS-*pum*^RNAi^*/+* Non-Deprived (N=31) and Deprived (N=32); UAS-*pum*^RNAi^*/tim-*Gal4 Non-Deprived (N=31) and Deprived (N=28). Because the calculations of sleep lost and sleep recovery involve both the Non-Deprived and Deprived groups (see methods), the N for panels A and B is equal to the N of the Deprived group. Data points and error bars represent means ± SEM. Stars indicate significance level (* denotes p<0.05; ** p< 0.01; *** p< 0.001; **** p< 0.0001).

So far, our findings link the duration of sleep deprivation to *pum* regulation, which is consistent with the expected role of neuronal homeostasis on sleep regulation. Since we observed greater homeostatic changes during chronic SD, we continued throughout the study using chronic SD to measure *pum*’s regulatory effects in compensatory sleep. The difference in sleep rebound between *pum*^RNAi^ vs parental flies does not seem to be related to non-specific effects of the genetic background affecting baseline sleep because daytime baseline sleep of *pum*^RNAi^ flies is higher than both parental and “sibling” controls (supplementary Fig S1). If baseline sleep would have been a contributing factor for the recovery results, we should have expected a higher sleep rebound. The fact that we obtained a lower rebound indicates pum knockdown rather that genetic differences influencing baseline sleep are the culprit of our results.

### *Pumilio* differentially changes expression level of genes associated with neuronal excitability in chronic vs acute SD

To determine if the reduction in homeostatic sleep rebound observed in *pum*^RNAi^ flies can be explained by changes in gene expression, we performed a quantitative reverse-transcription polymerase chain reaction (qRT-PCR) for a selected group of genes encoding synaptic proteins, synaptic translation modulators, neurotransmitter receptors and ion channels. In addition, we wanted to assess if the behavioral differences observed between acute vs chronic SD correlated with gene expression patterns. If Pum is necessary to reduce neuronal excitability caused by the high neural activity induced by SD, then knocking down *pum* should increase gene expression of synaptic proteins associated with neuronal excitability. In addition, if Pum recruitment is directly influenced by sleep need, as suggested by the behavioral differences between acute vs chronic sleep, then the increased sleep need during chronic SD would cause a differential expression of synaptic markers between acute and chronic SD.

For our analysis, we selected the synaptic genes *bruchpilot* (*brp*), *disks large 1* (*dlg1*) and *Synapsin* (S*yn*) as their protein products are known to increase after acute SD, as shown by western blots of whole fly brains (Gilestro, et al., 2009). In addition, we selected three genes that encode translation regulators —the *eukaryotic translation initiation factor 4E1* (*eIF4E1*), *Target of rapamycin* (*Tor*), and the Protein Kinase B (*Akt1*) because, as previously stated, EIF4E is a direct Pum target and both TOR and AKT are upstream regulators of EIF4E (Miron, et. al., 2003). We also included genes for the voltage gated sodium channel *paralytic (para)*, the voltage gated potassium channel *Shaker cognate l* (*Shal*) and *slowpoke* (*slo*), and the potassium channel modulator *sleepless* (*sss*, also known as *quiver* (*qvr*)), due to their relation to neuronal excitability. To complete the qRT-PCR testing panel, we also included the nicotinic Acetylcholine Receptor gene (*nAchRα1*), the GABA_A_ receptor gene *Resistant to dieldrin* (*Rdl*) and the *Glutamic acid decarboxylase 1* (*Gad1*), which synthetize for the enzyme that synthesizes the inhibitory neurotransmitter GABA (Lee, et al., 2003), because they also have been associated to regulations in neuronal excitability (see table S1 for references).

The RNA for the qRT-PCR study was extracted from whole heads, which were frozen two hours after the completion of the SD protocol. We evaluated the gene expression for non-deprived conditions against acute SD (12 hours) and chronic SD (84 hrs). The non-deprived results come from flies of each of the phenotypes handled in parallel to the deprived flies during the same experimental dates. First, we assesed the effects of *pum* knockdown within non-deprived flies on basal gene expression of our gene panel. Results show that the expression of *Shal* and *Gad1* was significantly increased in *pum*^RNAi^ flies as compared to the sibling controls (Fig. 3A). These results align with previous studies characterizing *pum* effects in neuronal excitability, which have shown a significant diminution of *Shal* mRNA when *pum* is overexpressed pan-neuronally (Murano, et. al., 2008). In addition, the expression increase in the inhibitory neurotransmitter synthesis enzyme *Gad1* was expected because Gad1 is a predicted target of Pum (Chen, et al., 2008). Furthermore, it has been shown that GABA acts as a slow inhibitory neurotransmitter in circadian neurons (Hamasaka, et al., 2005), promoting fly sleep (Parisky, et al., 2008). The fact that *pum*^RNAi^ flies showed increase levels of *Shal* and *Gad1* in non-deprived flies, suggests that the presence of Pum is also necessary to maintain normal sleep. This fact was corroborated by the increase in baseline sleep of *pum*^RNAi^ flies (supplementary Fig. S1), which should be expected under increased GABAergic inhibition of wake promoting neurons (Parisky, et al., 2008).

**Figure 3:**
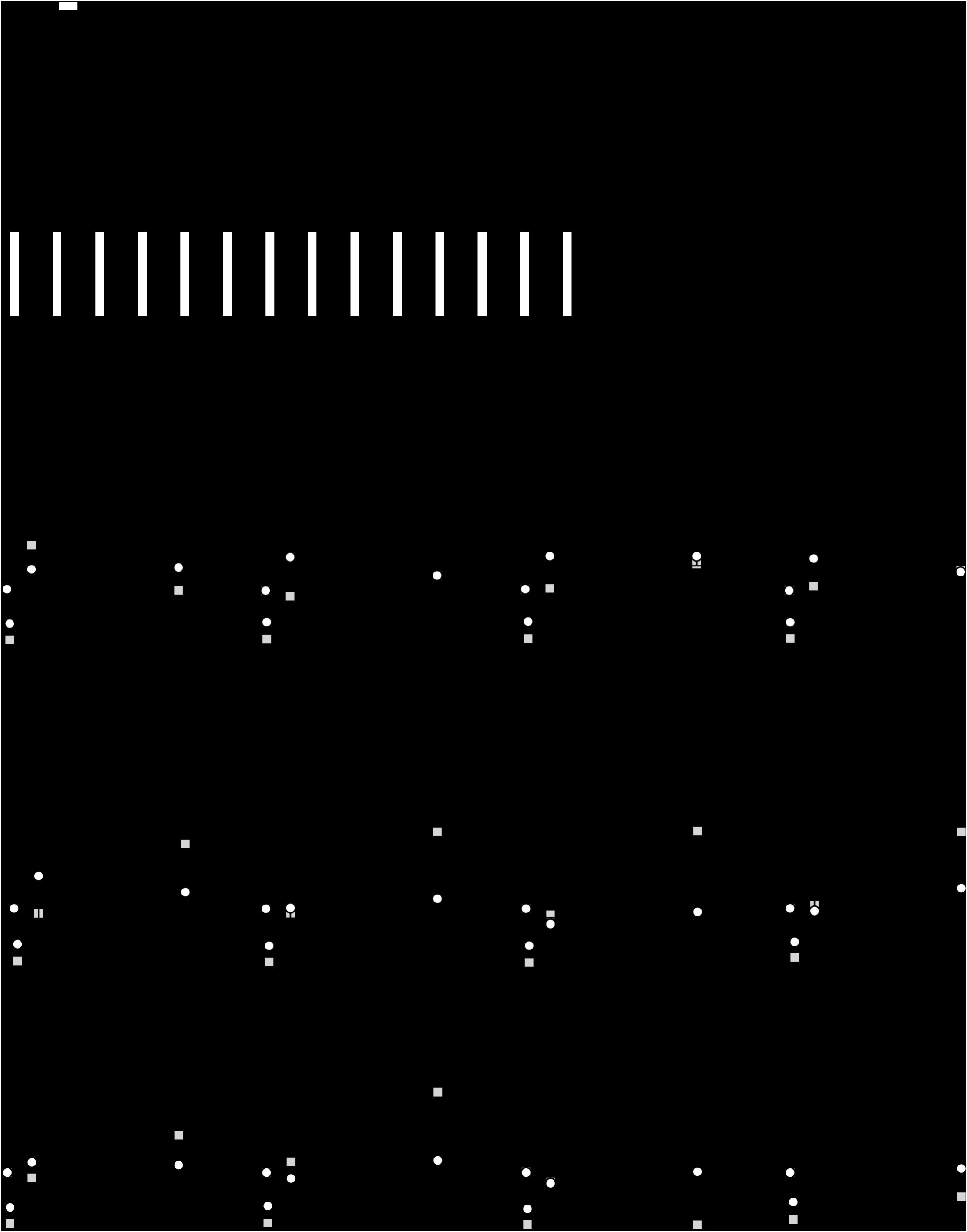
Pum knockdown results in differential expression patterns between acute (12 hours) and chronic (84 hours) sleep deprivation. Gene expression comparison of UAS-*pum*^RNAi^*/+* (“sibling” controls) vs UAS*-pum*^RNAi^*/tim*-Gal4 (experimental) subjected to acute (12 hours) mechanical SD vs chronic SD. **(A)** Baseline gene expression in non-deprived flies from both genotypes. **(B-E)** Time-course plots for non-deprived, acutely deprived and chronically deprived flies showing expression changes during acute deprivation. The fold change is expressed in log scale. **(F-J)** Time-course plots for non-deprived, acutely deprived and chronically deprived flies showing expression changes during chronic SD. The fold change is expressed in log scale. Data points and error bars represent means ± SEM. Two-way Analysis of variance (ANOVA) with repeated measures revealed significant effects due to *pum*, Time (T=0, T=12, T=84 hrs SD) and interactions between the parameters in some of the groups (see graphs for results). Stars indicate significance level (* denotes p<0.05; ** p< 0.01; *** p< 0.001; **** p< 0.0001).

Next, we assessed the changes in gene expression induced by acute and chronic SD, in both *pum*^RNAi^ flies and sibling controls. The qRT-PCR results showed that four genes displayed significant expression changes after acute SD but no change in response to chronic SD. These genes are: *nAchRα1, Rdl, para* and *slo* (Fig. 3B-E). For *nAchRα1*, this change was exacerbated in the *pum*^RNAi^ flies, whereas for *Rdl, para* and *slo*, the effect of acute SD in expression observed in control flies was abolished by the knockdown in *pum.* In contrast, eight different genes displayed significant changes between *pum*^RNAi^ flies and sibling controls in response to chronic SD, but no change in response to acute SD (Fig. 3F-M). A *pum* knockdown*-*dependent increase was observed in *eIF4E1, Tor, Akt, brp*, dl, and *Shal*; whereas a *pum* knockdown*-*dependent decreas was observed in *Syn* and *Gad1*. These results showed a concordance between the selected markers overexpressed by *pum’s* knockdown and their association with increased neuronal excitability. We observed gene expression increases in *pum*^RNAi^ flies but not in the “sibling” controls in synaptic translation genes like *eIF4E, Tor, Akt* (Penney, et al., 2012; Lee, et al., 2011; Howlett, et al., 2008) (Fig 3F-H); and genes coding for synaptic proteins like *brp* and *dlg* (Kittel, et al., 2006; Prange, et al., 2004) (Fig 3I-J). In addition, we saw an expression increase the *Shal* potassium channel (Fig 3K), which has been associated with neuronal excitability during repetitive locomotor activity (Ping, et al., 2011). We also saw an expression decrease in the synaptic protein gene *Syn* (Fig. 3L). The silencing of *Syn* increases intrinsic cell excitability associated with increased Ca^2+^ and Ca^2+^-dependent BK currents (Brenes, et al., 2015), which is also aligned with our expected results. In addition, *Gad1* was also less expressed in the *pum*^RNAi^ flies than their respective controls. These results are expected because GABAergic inhibition of wake promoting neurons has been shown to regulate sleep in *Drosophila* (Agosto, et al., 2008; Chung, et al., 2009). These combined results confirmed our hypothesis that *pum*’s effects in compensatory sleep behavior is correlated to changes in gene expression from selected neuronal excitability genes, and that acute vs chronic SD exhibit differential gene expression patterns, which points towards a differential regulation in acute vs chronic SD.

### *Pumilio* mutants show reduced sleep rebound

Finally, we used mutant fly lines to further validate our results independently of transgenic flies. To confirm the effects of *pum* knockdown in sleep homeostasis we selected the classical loss of function allele *pum*^13^ (also known as *pum*^*680*^). *Pum*^*13*^ is a dominant negative allele that bears a single amino acid substitution, which not only knocks down *pum* function but also interferes with normal *pum* function in heterozygotes (Wharton, et al., 1998). Thus, in addition to the semi-lethal *pum*^13^ homozygous mutants, we used *pum*^*13*^/*TM3* heterozygotes in our experiments.

The sleep deprivation produced similar sleep lost amounts in each of the lines tested. Fig 4A-C and D). Nonetheless, the sleep recovery showed a significant difference between both wild type (+/+) and *pum*^*13*^*/+* flies compared to *pum*^13^/*pum*^13^ flies (Fig 4E). By the end of the recovery period, the differences between *pum*^*13*^*/+* and the knockout *pum*^*13*^*/pum*^*13*^ were still maintained. Moreover, *pum*^13^/*pum*^13^ escaper flies completely abolished rebound to chronic sleep deprivation for the first 12 hours of the recovery period (Fig. 4E). This suggests that differential *pum* levels between the heterozygote and the *pum*^*13*^ homozygote, have correlative regulatory effects in sleep rebound.

**Figure 4:**
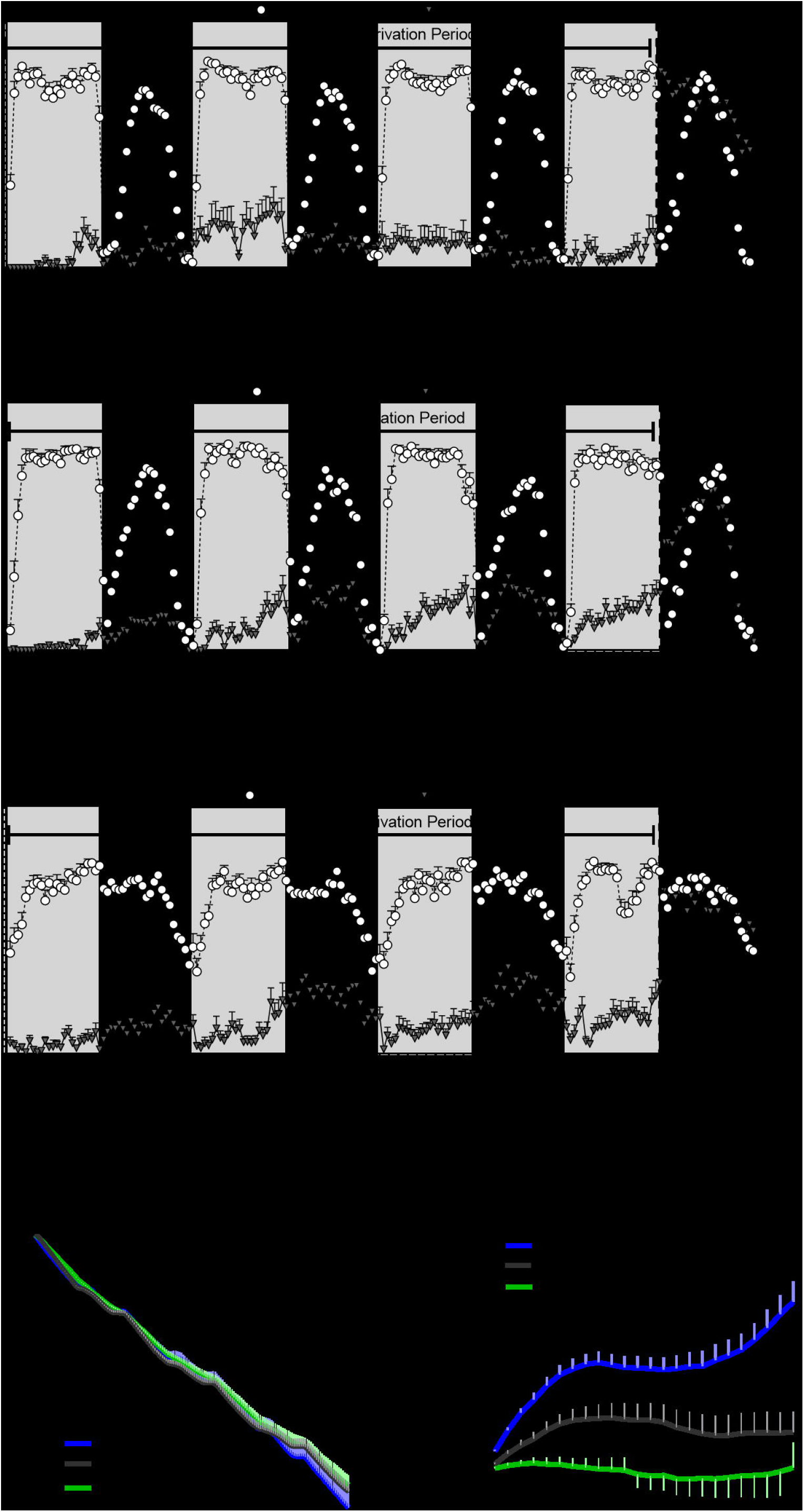
*Pum*^*13*^ mutant show reduced sleep rebound after chronic SD. Sleep comparison of wild type, heterozygous and homozygous flies for the *pum*^*13*^ allele. **(A-C)** Depiction of sleep activity during the sleep deprivation and sleep rebound period for all genotypes. The X axis indicates time after the start of the sleep deprivation protocol. The y-axis shows the number of minutes that flies slept in intervals of 30 min. **(D)** Cumulative sleep lost during deprivation expressed as a percentage of total sleep in non-deprived flies of the same genotype. Two-way ANOVA, using “genotype” as a factor and “time” as a repeated measure, did not show significant differences between the genotypes F (2, 63) = 0.3635, P=0.6967). **(E)** Percent sleep recovery after SD. Two-Way ANOVA with repeat measures indicated significant difference in genotypes (F (2, 63) = 11.29 P<0.0001) and interaction (F (46, 1449) = 5.667 P<0.0001). Post-hoc analysis using Uncorrected Fisher’s LSD comparisons test comparing all genotypes against *pum*^*13*^/*pum*^*13*^ flies revealed significant differences with *pum*^*13*^/+ flies (P=0.0319) and with *pum*^*13*^/*pum*^*13*^ (P<0.0001). The comparison between *pum*^*13*^/+ and *pum*^*13*^/*pum*^*13*^ show no difference (P=0.0728). The data shown represents one experiment with the following sample sizes (N): 1) Canton-S (+/+), Non-Deprived (N=30) and Deprived (N=17); *pum*^*13*^/+, Non-Deprived (N=28) and Deprived (N=28); *pum*^*13*^/*pum*^*13*^, Non-Deprived (N=30) and Deprived (N=22). Because the calculations of sleep lost and sleep recovery involve both the Non-Deprived and Deprived groups (see methods), the N for panels A and B is equal to the N of the Deprived group. Data points and error bars represent means ± SEM. Stars indicate significance level (* denotes p<0.05; ** p< 0.01; *** p< 0.001; **** p< 0.0001).

Additionally, we used the p-element insertion *pum* allele, Milord-1, to confirm the mutant results with another independent line. This line was generated by single transposon mutagenesis inserted in the *pum* transcriptional unit (Dubnau, et al., 2003). We compared this line with controls obtained from a wild type stock Canton S flies. As expected, Milord-1 flies showed a significant sleep rebound reduction (Fig 5D). Although there was a significant sleep lost difference between the genotypes at the end of the deprivation period (Fig. 5C), the ANOVA table results did not show a significant difference between the genotypes for the whole deprivation period. In addition, the sleep recovery calculation normalizes by the sleep lost, therefore, any sleep lost differences affecting the results have already been considered.

**Figure 5:**
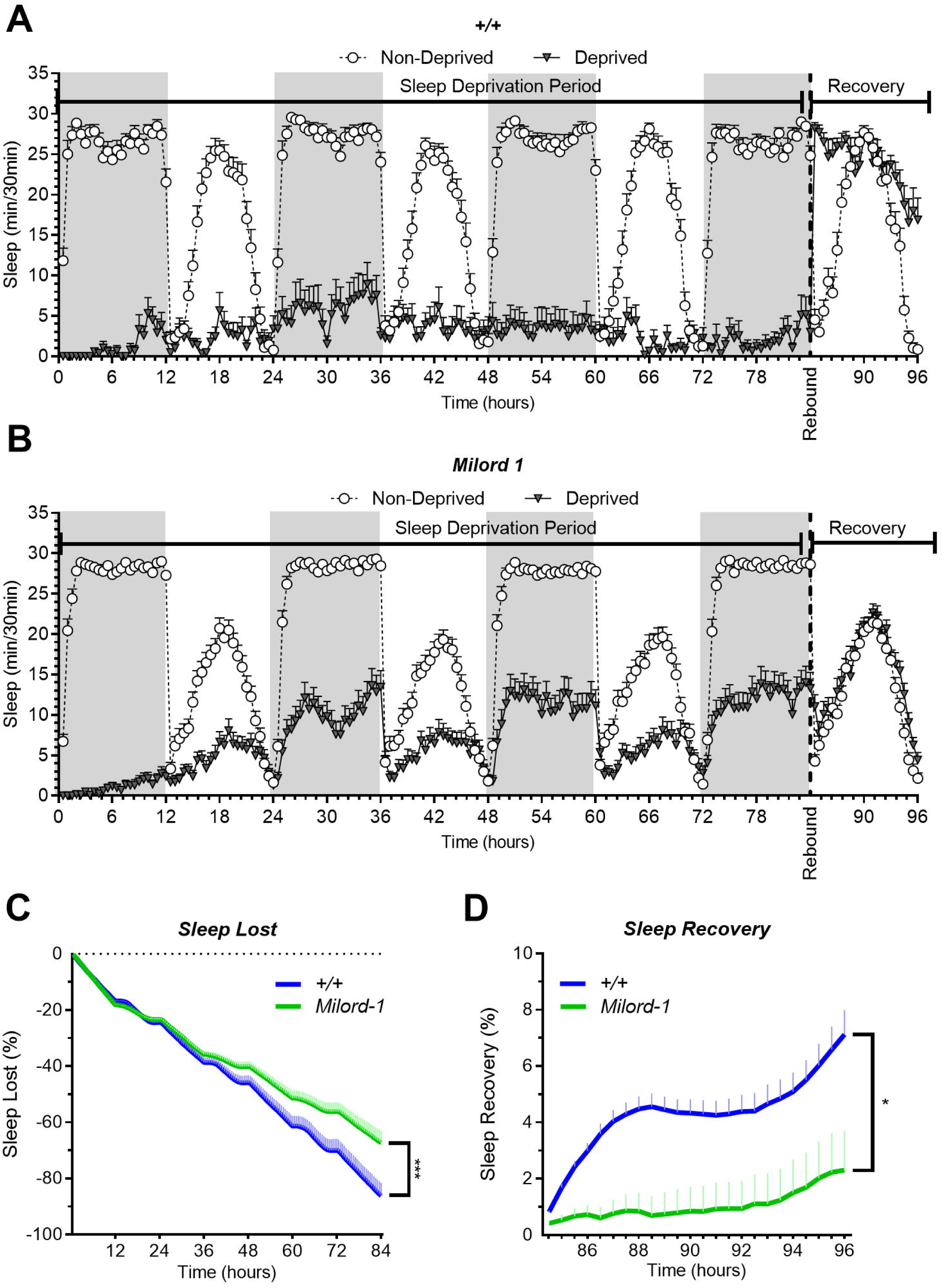
The Milord-1 fly line shows reduced sleep rebound after chronic SD. Sleep comparison of wild type and Milord-1 flies. **(A-B)** Depiction of sleep activity during the sleep deprivation and sleep rebound period for all genotypes. The X axis indicates time after the start of the sleep deprivation protocol. The y-axis shows the number of minutes that flies slept in intervals of 30 min. **(C)** Cumulative sleep lost during deprivation expressed as a percentage of total sleep in non-deprived flies of the same genotype. Two-way ANOVA using “genotype” as a factor and “time” as a repeated measure showed no significant differences between the genotypes (F (1, 58) = 3.712, P=0.0589). **(D)** Percent sleep recovery after SD. Two-Way ANOVA with repeat measures indicated significant difference in genotypes (F (1, 58) = 5.193 P=0.0264) and interaction (F (23, 1334) = 1.695 P<0.0213). The data shown represents two experiments with the following sample sizes (N): Canton-S (+/+) Non-Deprived (N=30) and Deprived (N=17); Milord-1 Non-Deprived (N=62) and Deprived (N=45). Because the calculations of sleep lost and sleep recovery involve both the Non-Deprived and Deprived groups (see methods), the N for panels A and B is equal to the N of the Deprived group. Data points and error bars represent means ± SEM. Stars indicate significance level (* denotes p<0.05; ** p< 0.01; *** p< 0.001; **** p< a 0.0001).

## Discussion

Through a combination of transgenic RNAi knockdown and mutant analysis, our results indicate that *pum* is necessary for the compensatory sleep behavior displayed after sleep deprivation in *Drosophila.* The *pum-*dependant regulation of sleep compensation, and its effects on synaptic gene expression, increases as sleep needs increases. Compensatory sleep rebound after a 12-hour sleep deprivation protocol (acute SD) was slightly reduced by knockdown of *pum* in *tim* neurons, but completely abolished after 84-hour of sleep deprivation (chronic SD). These differential effects were accompanied by a series of distinct changes in the expression of genes encoding synaptic proteins as well as synaptic translation factors. Together our data suggests that neuronal homeostasis mechanisms led by Pum differentially regulate compensatory sleep after acute and chronic SD, most likely through the regulation of neuronal excitability.

Interestingly, we also observed that *pum*^RNAi^ flies have increased day-time sleep in non-deprived conditions (Fig. 1, Fig. S1A), suggesting that other sleep behaviors are also regulated by *pum.* This effect of *pum* could perhaps be explained by the increased expression levels of *Gad1* and *Shal* in *pum*^RNAi^ non-deprived flies, as both genes are associated with a depression in overall neural activity. Additionally, the role of *pum* on regulating baseline sleep seems to be disconnected from its role in regulating sleep rebound. For instance, the daytime baseline sleep, in *pum*^RNAi^ flies is about two times the baseline of both control flies (Fig. S1A), but the same flies showed no rebound sleep after SD, suggesting that the homeostatic sleep rebound is independently regulated from baseline sleep. This interpretation is supported by reports from other groups. Shaw, et al, (2002) previously reported that *cycle* (*cyc01*) mutants showed an exaggerated response to sleep deprivation, which was 3 times as high as baseline sleep. In a similar way, Seidner, et al., (2002) found evidence suggesting that baseline sleep and homeostatic sleep can be regulated by distinct neural circuits.

Initial studies of chronic SD in other species have also pointed to a potential difference in the regulatory mechanisms between acute vs chronic SD. Rats exposed to chronic SD do not seem to regain the sleep lost even after a full 3-day recovery period, whereas in acute deprivation, most of the sleep was regained (Kim, et al., 2007). Critics attributed these differences, between acute and chronic SD, to the increase in sleep pressure, which force micro-sleep episodes or EEG artifacts during chronic SD (Leemburg, et al., 2010). A recent study showed that chronically sleep deprived animals no longer expressing the compensatory increases that characterize sleep homeostasis in daily sleep time and sleep intensity (Kim, et al., 2013). The authors of the study suggested that this decoupling of sleepiness from sleep time/intensity imply that there is one sleep regulation system mediating sleepiness (homeostatic), and another regulatory system for sleep time/intensity (allostatic) (Kim, et al., 2013). Whether the lack of sleep compensation observed during chronic SD is a real mechanistic phenomenon or an artifact of the deprivation method remained controversial. In our study, we wanted to test if the behavioral differences reported by the literature, between acute and chronic SD, were regulated by the same mechanism under the *pum* gene. Our results point to the presence of a differential homeostatic response between acute vs chronic SD in *pum* knockdowns, which suggests that *pum* participation in sleep homeostatic regulation is proportional to sleep need. Our data indicates that *pum* regulation of sleep rebound is done through the activation of different genes between acute and chronic SD. This difference seems to be aligned with fast action ion channel genes for acute SD and translation related and/or genes in which we expect to require more time to become active for chronic SD. Furthermore, we can hypothesize that individual neuroadaptations either promote or inhibit sleep rebound, and the neuroadaptations that promote rebound accumulate with sleep need. In this scenario, *pum* seems to be a key player among neuroadaptations promoting sleep rebound, which can be confirmed by the fact that *pum*^RNAi^ flies continued with a lower sleep recovery for a few days after SD was discontinued (Fig. 2S).

The qRT-PCR results support the hypothesis that *pum*^RNAi^ flies are in a higher excitable state than “sibling” controls. The significant expression increase observed in *nAchRα1* (Fig. 3B) during acute SD aligns with an increase excitability in *pum*^RNAi^ flies as acetylcholine is a major excitatory neurotransmitter. Furthermore, in mammals, acetylcholine has been shown to control the excitability of the circadian Suprachiasmatic nucleus (SCN) (Yang, et al., 2010). Also, *pum*^RNAi^ flies showed significantly less expression of the GABA receptor gene *rdl* compared to the “sibling” control (Fig. 3C). Previous studies have shown that reduced expression of *rdl* in PDF wake promoting neurons reduces sleep (Chung, et al., 2009), which could also explain the reduced sleep rebound of *pum*^RNAi^ flies. Additionally, the potassium channel *slo* also showed an increased expression in the “sibling” control compared to *pum*^RNAi^ flies. *slo* has been found to both increase or decrease neuronal excitability depending on the circuit where it was manipulated (Jepson, et al., 2013), therefore, we need to view this result in the context of the other gene expression changes.

The expression increases in *eif4e, Tor, Akt, brp, dlg*, and *Shal*, in *pum*^RNAi^ flies during chronic SD, are aligned with an expected increase in neuronal excitability induced by prolonged wakefulness and the knockdown of *pum* in the circadian circuit. Studies have shown that down-regulation of the Pum target eIF4E, reduced dendritic spine branching, thus affecting spine morphogenesis and synaptic function (Vessey, et al., 2010). Other studies have shown that TOR promotes retrograde compensatory enhancement in neurotransmitter release key to the homeostatic response in the *Drosophila* NMJ (Penney, et al., 2012). In addition, the levels of p-Akt increases strongly after glutamate application in *Drosophila* larvae (Howlett, et al., 2008). The *brp* mutants have shown impaired vesicle release and reduced Ca+ channels density in *Drosophila* neuro muscular junction (NMJ) (Kittel; et. Al., 2006), thus increased levels of BRP are important for efficient neurotransmitter release. In mice, the overexpression of Pum target Dlg (also known as PSD-95), resulted in enhanced excitatory synapse size and miniature frequency and a reduced the number of inhibitory synaptic contacts (Prange, et al., 2004). Moreover, blocking the potassium channel *Shal* in wake promoting neurons, delays sleep onset (Feng, et al., 2018), suggesting neuronal excitability of wake promoting neurons regulates sleep. Furthermore, *Syn*, which is associated with reserve vesicle release (Gitler, et al., 2008), showed a reduced expression in our qRT-PCR results. These results are also correlated to neuronal excitability. A study in mice reported increases in spontaneous and evoked activities in *Syn* knockouts (Chiappalone, et al., 2008). In sum, the expression changes of all these targets in sleep deprived UAS-*pum*^RNAi^/tim-Gal4 knockdown compared to the control flies demonstrates that the observed *pum* effects in chronic compensatory sleep can be associated with significant molecular changes aligned with changes into structural synaptic homeostasis that underlie an increased neuronal excitability in whole brain.

Out of the fourteen genes tested, only *para*, a direct Pum target, was contrary to our expectation during acute SD. Although *tim*-Gal4 is strongly expressed in glial cells (Kaneko & Hall, 2000), the circadian neurons expressing *tim*-Gal4 represent a relatively small number of cells in the fly brain, therefore, gene expression effects of *pum* knockdown over its direct molecular targets will be confounded with gene expression from the rest of brain cells. Nevertheless, it is reasonable to expect an indirect over-expression in a significant number of genes associated with neuronal excitability. Some of the relatively small number of circadian neurons in the fly brain have an important wake promoting role (Parisky, et al 2008), therefore they project widely into the brain and regulate a significant proportion of it. We hypothesize that knocking down *pum* in the circadian circuit avoids brain processes to “shut down” the neuronal excitability generated during chronic SD, hence the markers for increased neuronal excitability appear to be brain-wide over-expressed. It seems that prolonged sleep deprivation induces brain-wide changes in the expression of synaptic proteins and other neuromodulators, which trigger neuronal homeostatic processes to reduce neural activity. Our data supports the hypothesis that knocking down *pum* would disrupt this regulation allowing both the molecular expression and the behavioral activity of these flies to reflect a prolonged state of neuronal excitability.

The decrease in sleep rebound observed in *pum* knockdown is aligned with an increase in neuronal excitability, which was expected based on our hypothesis, by reducing the expression of the neuronal homeostasis gene *pum*. Pum is known to regulate sodium currents (Ina) and excitability in *Drosophila* motor neurons through translational repression and binding with *para*-RNA (Baines, et al., 2003), therefore reducing the number of available sodium channels. Reducing *pum* expression means there could be more sodium channels available and consequently, more neurons excited. Those excited neurons would have a diminished homeostatic mechanism to couple with the increased in excitability, resulting in prolonged wakefulness even after sleep deprivation stimulus was discontinued. Additional evidence in the literature supports the notion of a direct correlation between ion channels availability and wakefulness. Parisky, et al (2008), expressed the EKO potassium channel to hyperpolarize Ventral Lateral neurons (LNv) to reduce their excitability. In addition, they knocked down the *Shaw* potassium channel gene or expressed a dominant-negative Na+/K+-ATPase α subunit in the pdf LNv neurons in order to increase neuronal excitability. The results showed that suppressed LNvs increased sleep whereas hyperactive LNvs increased wake. Furthermore, studies in rats have shown increases in cortical neurons firing with increase in time awake (Vyazovskiy, et al., 2009). Moreover, Donlea, et al, (2014) found that the *crossveinless (cv-c)* mutants show decreased electrical activity in sleep promoting dorsal fan neurons. Additionally, the same study found that sleep pressure increases electrical excitability of sleep promoting neurons and this mechanism was blunted in *cv-c* mutants. This further strengthen our argument that *pum* regulates sleep homeostasis through the regulation of neuronal excitability. Identifying that a neuronal homeostasis gene, with a characterized mechanism of action, regulates sleep homeostasis, adds an important piece of information to further understand sleep homeostatic regulation.

Although this is the first time the neuronal homeostasis gene *pum* is linked to sleep homeostasis, there is additional evidence in the literature supporting the concept of neuronal homeostasis as a sleep regulatory mechanism. The neuronal homeostasis protein Homer mediates homeostatic scaling by evoking agonist-independent signaling of glutamate receptors (mGluRs) which scales down the expression of synaptic AMPA receptors (Hu, et al., 2010). Deletion of Homer in *Drosophila* produces fragmented sleep and failure to sustain long sleep bouts during sleep deprivation (Naidoo, et al., 2012). In addition, experiments where flies had a mutated shaker potassium (K+) channels exhibit reduced sleep (Cirelli, et al., 2005). The close functional relationship between neuronal sodium and potassium channels suggests the expression of sodium channels could also be associated with changes in the sleep phenotype. This was corroborated in experiments where a mutation in the sodium Na(v)1.6 channel gene, which *pum* regulates (Driscoll, et al., 2013), caused an increase in non-rapid eye movement (non-REM) sleep in rodents (Papale, et al., 2010).

Further studies characterizing additional Pum targets as well as other genes involved in neuronal homeostasis warrant exciting findings about the molecular control of sleep. Moreover, identifying the specific circuits where Pum is required for sleep regulation in both flies and mammals could provide a better picture of the mechanistic relationship between sleep function and molecular sleep regulation.

## Materials and methods

### Fly Stocks

Drosophila stocks were raised on standard Drosophila medium in a 12/12 h light/dark cycle. The following stocks were used in this study: The UAS*-pum*^RNAi^ (stock #26725: y[1] v[1]; P{y[+t7.7] v[+t1.8]=TRiP.JF02267}attP2) fly line was obtained from Bloomington Stock Center; The *tim*-Gal4 transgenic line: *yw; cyo/tim-*Gal4 was obtained from Dr. Leslie Griffith’s and Dr. Michael Rosbash’s labs at Brandeis University. These two lines were crossed to obtain both UAS-*pum*^RNAi^/*tim-*Gal4 experimental flies and the “sibling” control flies UAS-*pum*^RNAi^/+. The Milord-1 P{lacZ}^pummilord-1^ was obtained from Dr. Josh Dubnau. The mutant *pum*^*13*^ (*pum*^*680*^) and Canton S wild type flies were also obtained from Bloomington Stock Center and crossed to obtain both *pum*^*13*^*/+* and *pum*^13^/*pum*^13^ flies used in Figure 4.

### Sleep assays

Sleep assays used 1-2 days old female flies. The individuals were collected, separated by phenotype and placed into controlled temperature for 6-7 days under 12h:12h light dark cycles for entrainment. The individuals were then anesthetized with CO_2_ and placed in individual tubes containing fly food (5% sucrose, 2% agar). Tubes were then placed in *Drosophila* Activity Monitors (DAM) within an environmentally controlled incubator (26°C, 80% humidity, light intensity of 800 lux) and connected to the monitoring system (TriKinetics, Waltham, MA) under 12h:12h light dark cycles. After 4-5 days of baseline recordings, after changing the fly food to avoid dryness and microbial growth, the different groups of flies were sleep deprived with the methods described below. The genetic controls (“siblings”) were handled and tested side by side to the experimental flies. Flies with less than 80% deprivation within the first 12 hrs were excluded from the analysis. Number of individuals tested and number of experiment replications depicted are stated in figure legends. A cumulative sleep lost plot was calculated for each individual by comparing the percentage of sleep lost during sleep deprivation to the average sleep of the non-deprived flies. The individual sleep recovery (rebound) was calculated by dividing the cumulative amount of sleep regained by the total amount of sleep lost during deprivation.

### Mechanical sleep deprivation

Mechanical deprivation was performed using a commercially available *Drosophila* sleep deprivation apparatus (Trikinetics Inc., VMP Vortexer Mounting Plate). The apparatus was controlled by the Trikinetics software, shaking the monitors for 30 seconds on alternate settings of 4, 5 and 8 minutes to create an apparently random shaking pattern. The same pattern was used for all experiments. This set-up continued for 84 consecutive hours at the start of the first night for all chronic SD. For the acute SD experiment, the same set up was used but for only 12 hours of the deprivation night. Although this protocol results in partial sleep deprivation, rather than total deprivation, it induces significant sleep lost, normally around 80%, and allows the flies to survive through the chronic sleep deprivation period. Due to the long SD time of 84 hours and the baseline period, we perform a fly food change the day before SD to avoid microbial growth and food dryness. This change is coordinated with the morning peak and performed simultaneously for all experimental groups.

### Statistical methods

All statistical comparisons for significance between control and experimental groups was calculated using a significance cut off *p* < 0.05. All statistical analyses were performed using Graphpad Prism 8 software. Statistical analyses performed are included in the figure legends.

### Measurement of gene expression by qRT-PCR

RNA was extracted from heads of adult flies using the Qiagen RNeasy Mini kit (Qiagen, Crawley, UK). Five heads were pooled to make one sample and homogenized with a plastic mortar in 100ul of lysis buffer containing 0.1 M-mercaptoethanol, then 250 ul of lysis buffer was added and centrifuged. 350 ul of 70% ethanol was added and passed through a RNeasy column. After washing in buffer, immobilized nucleic acids were then treated with 190 U of DNase I for 15 min, washed again in stages according to manufacturer’s protocol, and then eluted in 20 ul of RNase-free water. Quantification of RNA concentration was made using a ND-1000 Nanodrop spectrophotometer (Nanodrop, Wilmington, DE). All extracted RNA samples were analyzed to assure quality using the Agilent Bioanalyzer, any samples showing signs of degradation were discarded. After adjusting for concentration, synthesis of cDNA was performed with the iScript Reverse transcription Supermix (Bio-Rad) as per manufacturer protocol. The mix was incubated at 25 °C for 5 min, then at 42 °C for 30 min followed by 85°C for 5 min to inactivate reverse transcription. From the total reaction volume of 20ul, 1 ul of cDNA was used for each PCR sample. All primers were obtained from Integrated DNA Technologies. An Eppendorf Mastercycler Thermal Cycler was used for the relative quantification of target mRNAs. Reactions contained 5 ul of Syber green (SYBR) (Invitrogen), 0.5 ul of each forward and reverse primer (both 10 mM), 3 ul of water, and 1 ul of cDNA. Cycling was as follows: initial denaturation of 15 sec at 95°C, then 40 cycles of annealing for 60 sec. for all primer pairs used, extension at 65°C for 1:20 min and melting curve generation at 95°C. Each group of 7 samples were tested in triplicate. Final mRNA levels were expressed as relative fold change normalized against *rp49* mRNA. The comparative cycle threshold (Ct) method (User Bulletin 2, 1997; Applied Biosystems, Foster City, CA) was used to analyze the data.

## Authors Contributions

J.L.A., N.R. and L.A.D. designed the study. J.A., N.R., C.J.P, J.O., R.N., M.F. and L.A.D. performed the experiments and data analysis. J.L.A, A.G.,N.F. and L.A.D. wrote/reviewed the manuscript.

## Acknowledgments

We thank Dr. Tugrul Giray, Dr. Adrienel Vazquez, Dr. Adrian Ovalos Dr. Manuel Giannioni, Hidequel Rodriguez, Bismark Madera, Gabriel Diaz, Maria del Mar Reyes, Franshesca Rivera, Alejandro Medina, Lizangelis Cueto, Melina Torres, Carlos Billini, Rosa Alers, Wilfredo Soto, Rubielis Serrano, Keila Velazquez, Marcelo Francia, Oto Mendez, Norelis Diaz and the students from the genetics lab at UPR-RP biol3350 for their support.

## Supplementary Information

**Figure S1: Transgenic flies showed increased baseline sleep. (A)** Graphs showing the average sleep activity for UAS*-pum*^RNAi^*/+* (“sibling” control) and UAS*-pum*^RNAi^*/tim-*Gal4 under baseline sleep conditions compared to parental *tim-*Gal4*/+* baseline. The y-axis shows the number of minutes that flies slept in intervals of 30 min. **(B)** Graph showing baseline sleep for all *pum*^*13*^ lines. **(C)** Graph showing baseline sleep for Milord-1 line. Stars indicate significance level (* denotes p<0.05; ** p< 0.01; *** p< 0.001; **** p< 0.0001).

**Figure S2: *Pum* knockdown shows reduced sleep recovery up to 96 hours after chronic sleep deprivation.** Sleep comparison of UAS*-pum*^RNAi^*/tim-*Gal4 (experimental) vs UAS-*pum*^RNAi^*/+* (“sibling” controls) for the period following chronic SD. **(A-B)** Depiction of sleep activity during the recovery period for both genotypes after chronic mechanical SD. **(C)** Extended percent sleep recovery after SD. Graph depicting up to 108 hours of sleep recovery after chronic SD. Two-Way ANOVA with repeat measures indicated significant differences between the genotypes (F (1, 80) = 18.1 P<0.0001) and interaction (F (167, 13360) = 8.362 P<0.0001). Post-hoc analysis using Tukey’s multiple comparisons test revealed significant differences between UAS*-pum*^RNAi^*/tim-*Gal4 throughout the recovery period. The y-axis shows the number of minutes that flies slept in intervals of 30min. The data shown represents two experiments with the following sample sizes (N): UAS-*pum*^RNAi^*/+* Non-Deprived (N=60) and Deprived (N=39); UAS-*pum*^RNAi^*/tim-*Gal4 Non-Deprived (N=63) and Deprived (N=43). Because the calculations of sleep lost and sleep recovery involve both the Non-Deprived and Deprived groups (see methods), the N for panels A and B is equal to the N of the Deprived group. The y-axis shows the number of minutes that flies slept in intervals of 30min. Stars indicate significance level (* denotes p<0.05; ** p< 0.01; *** p< 0.001; **** p< 0.0001.

**Figure S3: *Pum***^**RNAi**^ **acute SD time course from qRT-PCR flies confirmed acute SD differences in sleep rebound** Sleep comparison of *UAS-pum*^RNAi^*/tim-*Gal4 (experimental) vs UAS-*pum*^RNAi^*/+* (“sibling” controls) during acute and chronic SD. Flies were removed from the monitors after two hours of sleep recovery and immediately freeze for qRT-PCR analysis. **(A**,**C)** Depiction of the acute sleep deprivation and sleep rebound period for both genotypes. The y-axis shows the number of minutes that flies slept in intervals of 30 min. The data shown represents one experiment with the following sample sizes (N): UAS-*pum*^RNAi^*/+* Non-Deprived (N=31) and Deprived (N=27); UAS-*pum*^RNAi^*/tim-*Gal4 Non-Deprived (N=31) and Deprived (N=32). **(B**,**D)** Depiction of the sleep deprivation period and sleep rebound pattern for *tim-*Gal4*/+* (parental) flies, UAS*-pum*^RNAi^*/+* (sibling) and UAS*-pum*^RNAi^*/tim-*Gal4 exposed to chronic (84hrs) mechanical SD. The data shown represents two experiments with the following sample sizes (N): UAS-*pum*^RNAi^*/+* Non-Deprived (N=62) and Deprived (N=34); UAS-*pum*^RNAi^*/tim-*Gal4 Non-Deprived (N=61) and Deprived (N=54). Error bars represent means ± SEM.

**Table S1**: Summary of PR-PCR results in relation to each marker’s effect in neuronal excitability.

