## Supplemental Table 1 for "*pumilio* regulates sleep homeostasis in response to chronic sleep deprivation in *Drosophila melanogaster*"

| Table S1: Summary of qRT-PCR results in relation to each marker's effect in neuronal excitability | | | | | |
|---|---|---|---|---|---|
| **Gene** | **Gene Name** | **Results** | **Description** | **Relation to neuronal activity** | **References** |
| ***brp*** | *bruchpilot* | Over-expressed due to *pum* effect | Scaffolding synaptic protein | Increase amplitude of miniature excitatory junctional currents (mEJCs) | Kittel, et al., 2006 |
| ***dlg1*** | *discs large 1* (PSD-95) | Over-expressed due to *pum* effect | Scaffolding synaptic protein | Enhances excitatory synapse size and miniature frequency | Prange, et al., 2004 |
| ***Syn*** | *Synapsin* | Under-expressed due to *pum* effect | Synaptic protein | Knockdown increases spontaneous and evoked activities | Chiappalone, et al., 2009 |
| ***Csp*** | *Cysteine string protein* | No change | Synaptic protein | Promotes neuronal homeostasis | Brusich, et al., 2015 |
| ***eIF4E1*** | *eukaryotic translation initiation factor 4E1* | Over-expressed due to *pum* effect | Translational regulation | Promotes retrograde compensatory enhancement in neurotransmitter release | Penney, et al., 2012 |
| ***Tor*** | *Drosophila Target of rapamycin* | Over-expressed due to *pum* effect | Translational regulation | Promotes retrograde compensatory enhancement in neurotransmitter release | Penney, et al., 2012 |
| ***Akt1*** | *Akt1* (Protein Kinase B) | Over-expressed due to *pum* effect | Serine/Threonine Kinase | Knockdown prevented the insulin-induced increase in dendritic spine density | Lee, et al., 2011 |
| ***sss*** | *quiver* (sleepless) | No change | Ion channel regulator | Decreases neuronal excitability by antagonizing nicotinic acetylcholine receptors (nAChRs) | Wu, et al., 2014 |
| ***Para*** | *paralytic* | No change in chronic SD<br><br>Under-expressed in Acute SD $pum^{RNAi}$ | Ion channel (Na(v)) | Increase neuronal excitability | Mee, et al., 2004 |
| ***Shal*** | *Shaker cognate l* | Over-expressed due to *pum* effect | Ion channel (K) | Increases neuronal excitability when mutated, reduces excitability when open. | Parrish, et al., 2014<br><br>Ottosson, et al., 2015 |
| ***Slo*** | *slowpoke* | No change in chronic SD<br><br>Under-expressed in Acute SD in $pum^{RNAi}$ | Ion channel (BK) | Increases neuronal excitability by shortening refractory period<br>Effects in neuronal excitability are circuit dependent | Ghezzi & Atkinson, 2011<br><br>Jepson, et al., 2013 |
| ***nAChRα1*** | *nicotinic Acetylcholine Receptor* | No change | Ion channel | Increases neuronal excitability | Wu, et al., 2014 |
| ***Rdl*** | *resistant to dieldrin* | No change in chronic SD<br>Under-expressed Acute SD in $pum^{RNAi}$ | Ion channel ($GABA_A$ ) | Decreases neuronal excitability | Parisky, et al., 2008 |
| ***Gad1*** | *Glutamic acid decarboxylase 1* | Over-expressed due to *pum* effect<br>Under-expressed after Chronic SD due to *pum* effect | $GABA_A$ synthetase | Chemical increase in GABA rescued larvae from lethality associated with reduced excitability | Li, et al., 2014 |
