## Supplementary figures and images for "*pumilio* regulates sleep homeostasis in response to chronic sleep deprivation in *Drosophila melanogaster*"

### Supplemental Figure S1

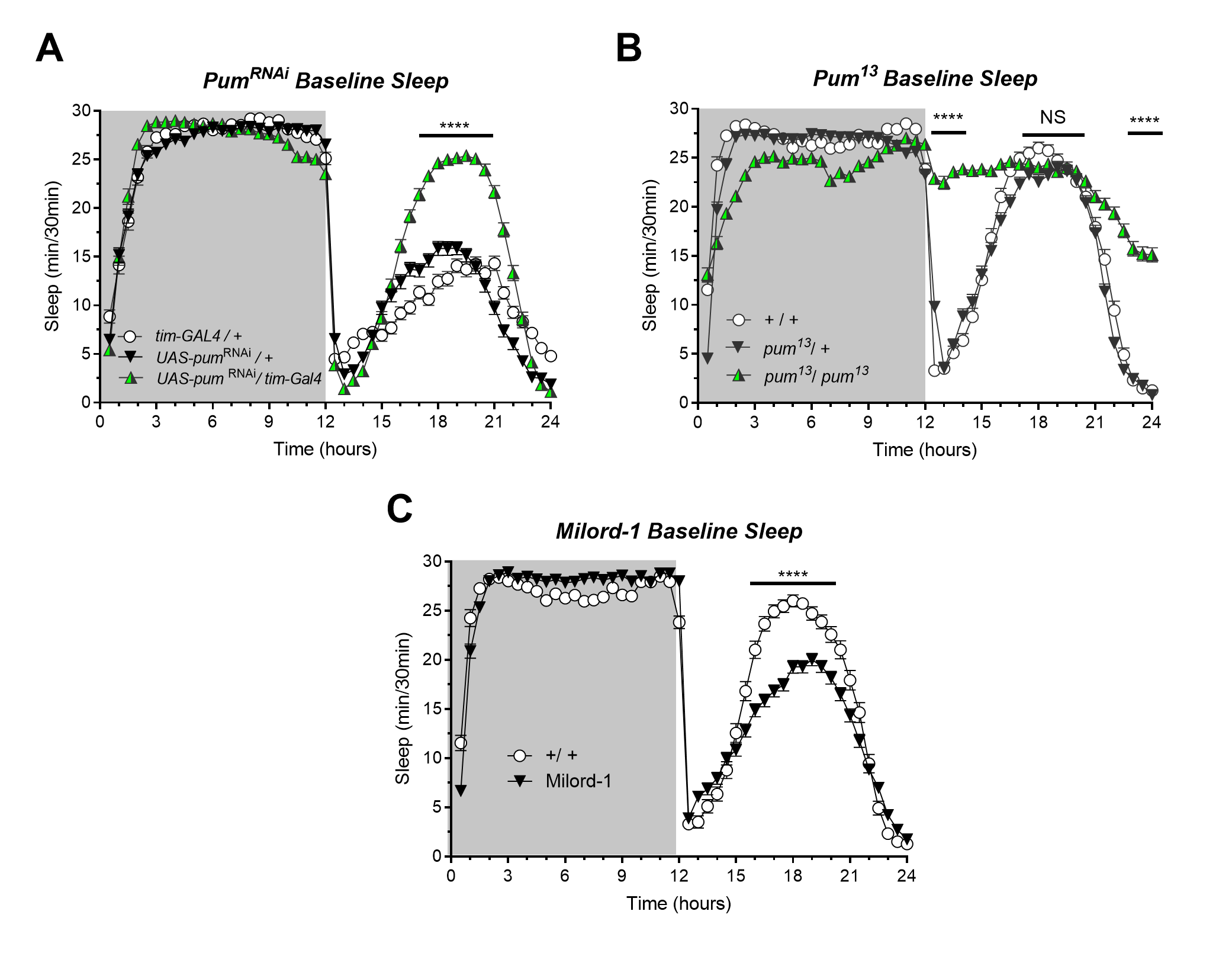

### Supplemental Figure S2

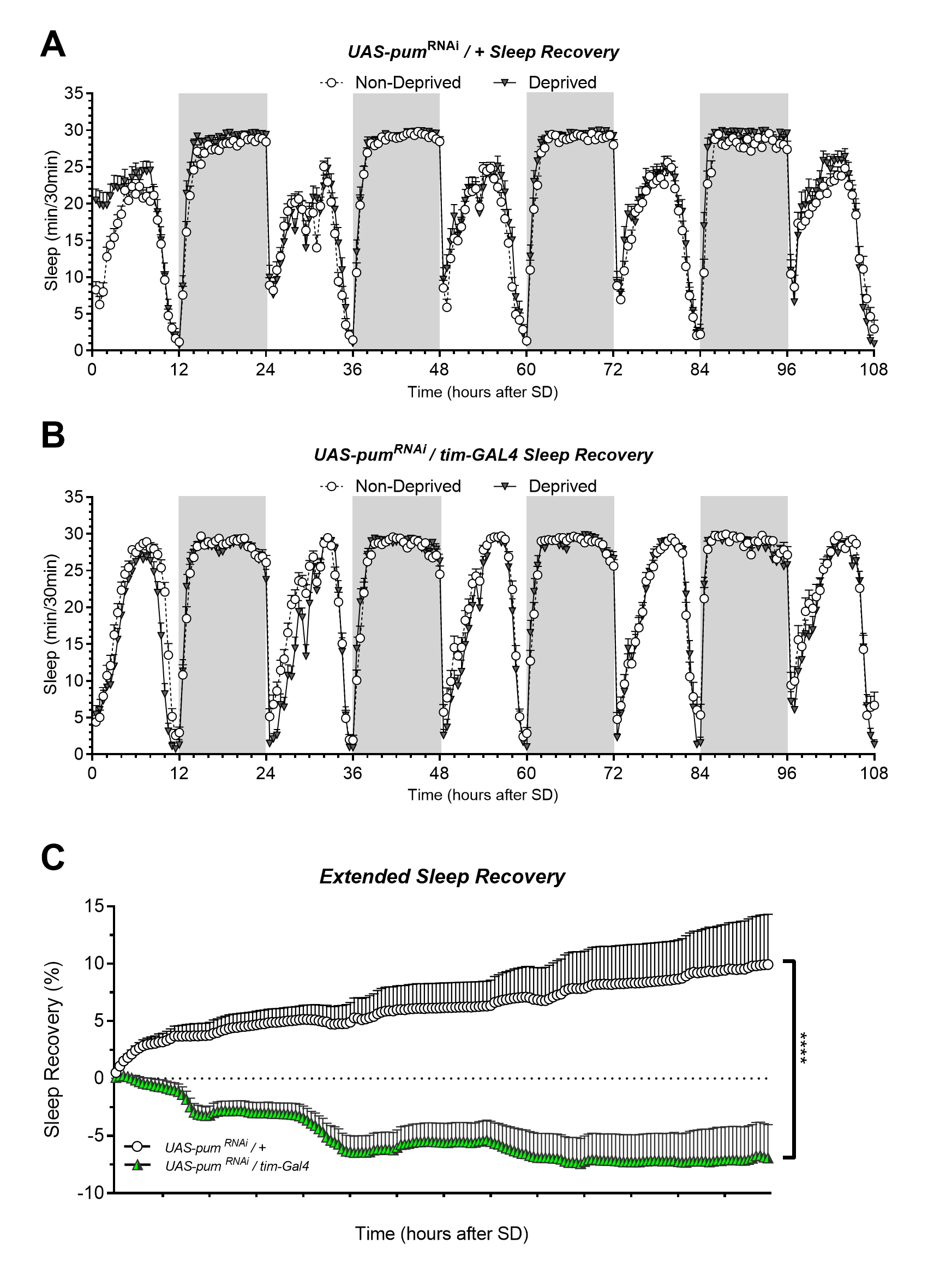

### Supplemental Figure S3

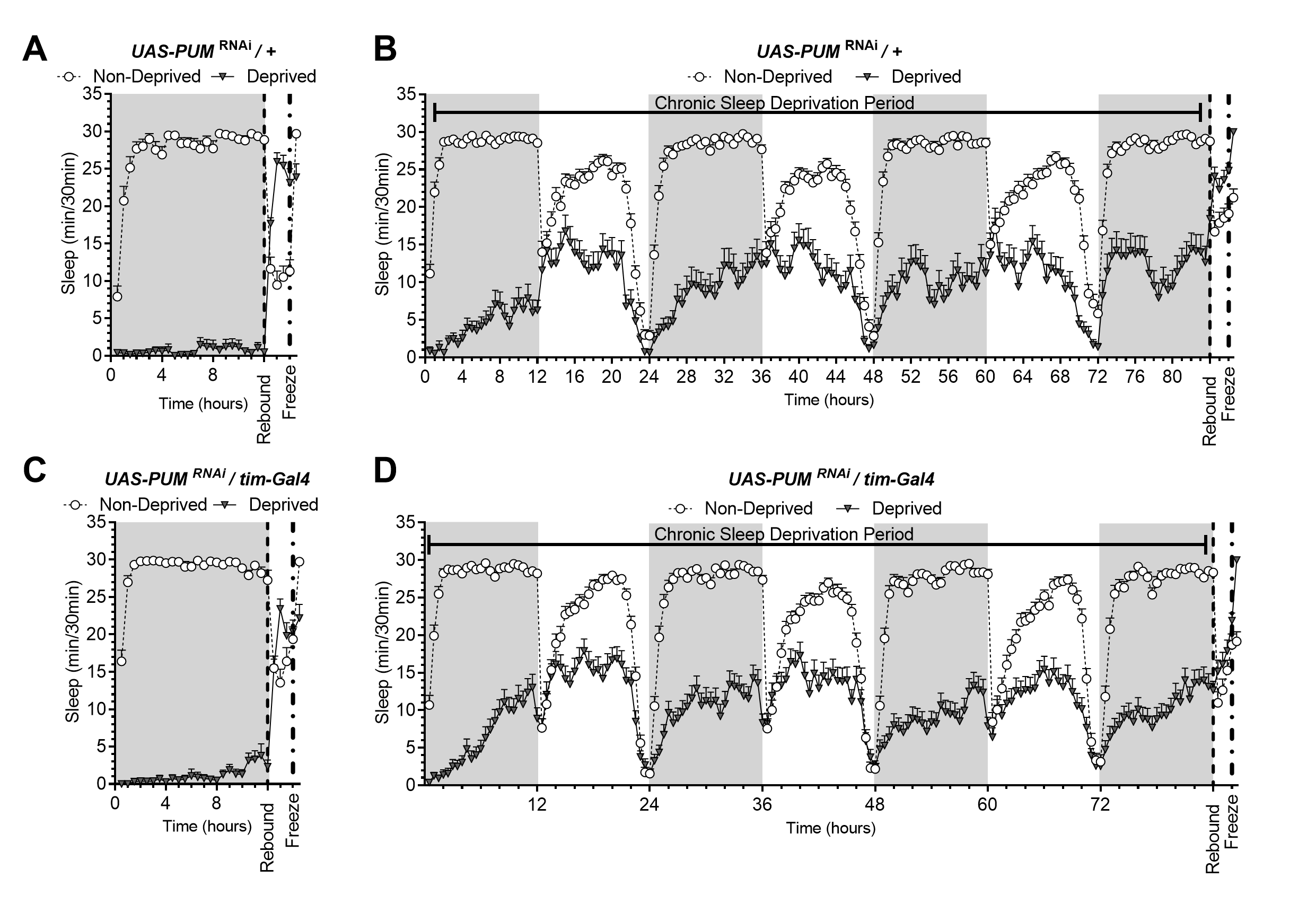
